# Spatial and temporal drivers of genomic inbreeding in a declining population of yellow-bellied marmots

**DOI:** 10.64898/2026.11.23.768605

**Authors:** Bridgett M. vonHoldt, Stavi R. Tennenbaum, Rose M. Nicholson, Michelle White, Yara M. Almoussa, Aiyonne F. Bryant, Anna Carvajal, Marina B. Cookson, Jamil Fayad, Julia Felice, Hannah Floyd, Sarah Geerges, Leila M. L. Granier, Navneeth Gurachar, Hayden P. L. Hajovsky, Mulin Huan, Cevina Hwang, Sophie R. Johnson, Alexandra Kelly, Wee-Ern Lau, Bethany Mariel Suliguin, Brianna Melanie Suliguin, Aya Surheyao, Kathleen Y. Tang, Daniel T. Blumstein, Julien G. A. Martin

## Abstract

Estimates of genetic diversity and effective population size are critical for predicting the resilience of natural populations to environmental change. We generated the first estimates of genetic diversity, effective population size and genomic inbreeding in a wild population of yellow-bellied marmots (*Marmota flaviventer*) studied continuously in western Colorado since 1962. Using 21 years of data from 284 individuals genotyped at 153,550 minimally filtered SNPs, including 17,393 neutral and unlinked SNPs, we found cryptic patterns of genetic subdivision and migration along a latitudinal gradient. A geographic cline in ancestry revealed a narrow (∼900 m) genetic contact zone consistent with limited, unidirectional gene flow from south to north on the landscape. We also discovered a recent, severe decline in both census size and Ne following the winter of 2010 from which the population has yet to recover, contrasting prior trends of demographic expansion over the past century. Genomic inbreeding has remained elevated since this population decline, although long runs of homozygosity remain low in frequency despite pedigreed mating between relatives. Higher maternal inbreeding was associated with lower seasonal mass gain in offspring, consistent with intergenerational inbreeding depression in a trait predictive of annual survival. These findings indicate that while this well-monitored population has persisted at a low Ne with relatively high inbreeding, a recent, indeterminate census collapse and evidence for inbreeding depression in a key fitness-linked trait may threaten its future persistence.

## Introduction

Anthropogenic stressors have led to shrinking habitat and population sizes across diverse taxa, reducing genetic diversity globally (Exposito-Alonso et al., 2022). Extensive theoretical and empirical work has shown that the adaptive capacity for a population to respond to environmental changes deteriorates as genetic diversity declines and populations become small and isolated (Allendorf, 2017; Frankham, 2005; Hoffmann et al., 2017; López-Cortegano et al., 2019). With these coupled threats to biodiversity, precise estimates of effective population size (Ne) and inbreeding are critical to estimate risks of inbreeding depression in a given population (vonHoldt et al., 2024), i.e., the reduction in fitness from consanguineous mating (Keller & Waller, 2002; Huisman et al., 2016). Inbreeding depression can erode individual and reproductive fitness by enriching recessive deleterious variants genome-wide (Charlesworth & Charlesworth, 1987; Charlesworth & Charlesworth, 1998), yet its extent can be difficult to quantify across organismal traits and life stages (Charlesworth & Willis, 2009). One reliable metric of inbreeding depression is the genomic quantity and length of runs of homozygosity (ROH), and correspondingly, the proportion of an individual’s autosomal genome that is covered by runs of homozygosity (FROH) (Kyriazis, Robinson & Lohmueller, 2025). These ROH segments indicate continuous stretches of DNA where two alleles at a locus are identical by descent (IBD). Long ROHs are indicative of recent inbreeding, low Ne or bottlenecks, and can be used to estimate deleterious genetic load (Kyriazis et al., 2023). More recently, ROHs have been used to directly quantify the loss of fitness from recent inbreeding (Szpiech et al. 2013) and to model future extinction risk (Kyriazis et al., 2026).

The pedigree-based inbreeding statistic (*F*) can also estimate the probability that two alleles are IBD, yet this metric relies on comprehensive breeding records that are often incomplete or unobtainable for all individuals beyond ex situ populations (Pemberton, 2008; Kardos et al., 2015). Pedigree inbreeding estimates have been supplemented using microsatellites (Huisman, 2017), but these approaches are prone to underestimate inbreeding due to limited marker resolution (Balloux et al., 2004) and accuracy loss from drift (Putman & Carbone, 2014). Advances in low-cost, low-coverage genome sequencing can now be leveraged to resolve kinship and estimate recent and historical inbreeding patterns at thousands of single nucleotide polymorphisms (SNPs) (Xia et al., 2016). Genomic inbreeding estimates using SNP data are powerful tools for characterizing the extent and fitness consequences of genetic diversity loss in wildlife over spatial and temporal scales, particularly useful in far-dispersing, difficult to observe or endangered species with complex mating structure (Kardos et al., 2015; vonHoldt et al., 2024a). Genomic data can also reveal natural drivers of elevated homozygosity, including dispersal barriers, reproductive skew, social dominance hierarchies and age (Cooney & Bennett, 2000; Cohas et al., 2009; Costello et al., 2008; Nelson-Flower et al., 2018; Li & Kokko, 2019; vonHoldt et al., 2024b; Loera et al., 2026).

Yellow-bellied marmots (*Marmota flaviventer*) are semi-social polygamous rodents found at high elevations across western North America (Olson & Blumstein, 2010; Blumstein, 2025). This species and others in *Marmota* show strong female philopatry (Allaine, 2000) while males tend to disperse (Armitage, 2012; Blumstein et al., 2013; Armenta et al., 2019), resulting in familial groups of related females at geographic colonies (Blumstein, 2025) that range from consistently to intermittently inhabited (Blumstein et al., 2009). Prior studies in this genus have found persistent low levels of genetic diversity linked to Pleistocene climate change, local adaptation to cold tolerance and long-term isolation of these lineages (Floyd et al., 2005; Bai et al., 2019; Gossmann et al., 2019). Despite the broad continental range of yellow-bellied marmots compared to most European species (Rankin et al., 2019), several *Marmota* species endemic to North America are in decline or critically endangered despite historically large census sizes (Kerhoulas et al., 2015; Frey et al., 2019), suggesting these cold-adapted species may be vulnerable climate change in montane ecosystems, possibly compounded by inbreeding depression within small, isolated subpopulations.

Here, we assessed genetic diversity and inbreeding depression in a subpopulation of yellow-bellied marmots (hereafter, “marmots”) on the western slope of the Colorado Rocky Mountains. This population has seen contrasting fitness responses to environmental change over recent decades: snowpack loss and warming over the last 40 years have decreased adult annual survival but increased juvenile recruitment (Cordes et al., 2020; Wells et al., 2020). We hypothesized that genomic inbreeding has increased due to combined effects of climate change and anthropogenic encroachment, which have impacted a suite of heritable life history traits (Blumstein, 2025; Adler et al., 2026). All prior genetic work in this population has relied on behavioral observations and pedigree construction using 10-12 microsatellite markers (Olson et al., 2010), which indicated that marmots do not avoid inbreeding despite apparent fitness costs (Olson et al., 2012). A lack of any genomic work and noted changes in census size (Ozgul et al., 2010) suggests that genetic diversity and inbreeding may have change over the study’s history. We further hypothesized that fine-scale geographic isolation restricts gene flow beyond earlier estimates. Evidence suggests coarse isolation-by-distance effects in marmots. Numerous local studies have identified divergent fitness outcomes and behavioral variation between resident colonies based on geography, and many have hypothesized that the effects of snowmelt timing, seasonal resource abundance, dispersal and social hierarchies may limit mating between colonies (Blumstein et al., 2013; Maldonado-Chaparro et al., 2015; Dumas et al., 2026). Despite these findings, the role of geographic variation in structuring genetic variation, gene flow and inbreeding in this species remains unknown.

We leveraged two recent decades of genomic data from individual marmots followed at natal colonies from birth into adulthood to first test whether geographic isolation maintains genetic structure. We then combined genomic inbreeding estimates with pedigree reconstruction to test if inbreeding levels restrict mass gain, the primary predictor of overwinter survival in marmots. We analyzed genetic diversity, ancestry, relatedness and inbreeding over fine spatial and temporal breadths at this site to evaluate potential risks to the stability and genetic health of this population in the future. As other montane and subalpine species face novel anthropogenic pressures with climate change, understanding how genetic diversity and inbreeding interact to influence demographic trajectories prior to conservation concern is critical to determine which natural populations may become threatened by genomic erosion.

## Materials and Methods

### Field sampling and DNA extraction

We collected blood and buccal swab samples from marmots near the Rocky Mountain Biological Laboratory (RMBL), Colorado, USA (38°57′ N, 106°59′ W) at nine study colonies (between 2,700-3,100 meters) during their active season from mid-April until early September. We live trapped marmots using Tomahawk (WI, USA) traps between 2004-2025 and restrained animals by hand or in canvas bags to collect 3 mL of whole blood in EDTA vacutainers (University of California, Los Angeles IACUC protocol 2001-191-01; RMBL and Colorado Parks and Wildlife TR917). We also collected buccal cells by thoroughly swabbing the inside of each cheek with the ORAcollect-Dx OCD-100 DNA collection kit (DNA Genotek, CA). We tagged each marmot with two unique numeric ear tags and marked their dorsal pelage with black Nyanzol fur dye for visual identification (Blumstein, 2013). We weighed captured animals using digital scales and recorded the sex and resident colony of each animal (>0.9 recapture rates at long-term study sites) (Ozgul et al., 2006; Ozgul et al., 2007). For juveniles (hereafter, “pups”), we recorded their year of birth (YOB), date of emergence from natal burrows (usually in June; Armitage & Downhower, 1974) and their body mass on emergence (in grams). We extracted genomic DNA (gDNA) from whole blood and buffy coat samples using the DNeasy Blood and Tissue Kit following the standard manufacturer protocol (Qiagen, DE). To isolate gDNA from ORAcollect-Dx OCD-100 collection swabs, we incubated the collection tubes for 24 hours at 50° C. We then conducted a bead-based buffer exchange and purification with SPRIselect paramagnetic beads (Beckman Coulter, USA) and eluted gDNA in Tris-HCl pH 8 (10mM). We quantified gDNA with the Quant-iT PicoGreen dsDNA Assay (Invitrogen, USA).

### DNA library preparation

We prepared purified gDNA for restriction site-associated DNA library sequencing (RADseq), described in further detail in (Ali et al., 2015). Briefly, we digested 75ng of gDNA with the Sfb1-HF restriction enzyme (New England Biolabs, USA), to which we ligated a unique 8bp biotinylated BestRAD adapter to each library before pooling multiple libraries for downstream preparation. We used SPRIselect beads for all library purification and size selection steps. First, we fragmented the pooled libraries to ∼300-400 bp on a LE220 Covaris Focused Ultrasonicator at the Princeton University Lewis Sigler Genomics Core. We enriched sheared library pools using Streptavidin Dynabeads M-280 (Invitrogen, USA) to capture adapter-ligated fragments that contained the biotinylated adaptors. Next, we used the NEBNext Ultra II DNA Library Prep Kit (New England Biolabs) to repair fragment ends, ligate the Illumina P2 i5 adapter to incorporate a second unique barcode to each pooled library and amplify target gDNA fragments using 14 cycles of PCR. We collected a minimum of 4 million reads per sample (2 x 150 nt) on the Illumina NovaSeq 6000 platform at the Yale University Center for Genome Analysis, New Haven, CT.

### Bioinformatic processing

To demultiplex the resulting sequences per individual from each pool, and to find and genotype polymorphic SNP loci, we used *STACKS* v2.6 (Catchen et al., 2013; Rochette et al., 2019). We used the *process_radtags* module to retain reads that carried ≤2 nucleotide mismatches to the expected barcode sequence and had a Phred quality score ≥10. We then used the *clone_filter* module to remove duplicated reads, before mapping to the yellow-bellied marmot genome (GenBank Accession GCF_003676075.3, SRA BioProject PRJNA516936) using *bwa-mem* v0.7.19 (Li, 2013). We converted files to the BAM format with *samtools* v0.1.18 (Li et al., 2009) and merged all BAM files that came from the same individual. We discovered and built a catalogue of polymorphic variants (SNPs) using the Marukilow model in the *gstacks* and *populations* modules in *STACKS* (--vt-alpha and --gt-alpha with *p*=0.01). We then used multiple filtering thresholds to construct three different SNP datasets of high-confidence genotyped loci. First, we used *VCFtools* v0.1.17 (Danecek et al., 2011) to exclude loci that carried singleton or private doubleton alleles, had <10-fold sequencing coverage, were missing a genotype in >40% of the samples, had a minor allele frequency (MAF) <0.03, and had >10% missingness per locus (-geno 10) in *PLINK* v1.9 (Chang et al., 2015). We refer to this as our “minimally filtered” SNP dataset.

### Genome-wide estimates of autozygosity

We used our minimally filtered dataset for analyses which relied on genetic linkage information and SNP density, such as estimating genomic autozygosity. For this metric of inbreeding, we further filtered any individuals with >20% per-sample missingness, before estimating each individual’s genomic runs of homozygosity (ROH) using *roh* in *bcftools* (--AF-dflt 0.5 -G 30) (Narasimhan et al., 2016). We retained only runs that had a Phred quality score ≥30. We then converted individual ROH values to the genomic inbreeding coefficient (FROH) by dividing the summation of the total length of ROH by the total length of the marmot genome (2,277,643,183 bp), defined as the distance between the first and last genotype per contig (McQuillan et al., 2008). We calculated the number, mean length and size distribution of ROH segments per individual. We used Wilcoxon rank-sum tests in R (R Core Team) to analyze statistical differences in FROH within the population by sex, colony, sampling year and an animal’s YOB.

### Genetic substructure and differentiation

To assess genetic structure, we constructed a statistically neutral and unlinked “demographic” dataset by removing loci in linkage disequilibrium (LD) and redundant, LD-correlated SNPs (r^2^>0.1) across 50 SNP windows 5 variants at a time, in *PLINK* (--indep-pairwise 50 5 0.1) and SNPs that deviated significantly from Hardy-Weinberg Equilibrium (--hwe, *p*=0.001) (vonHoldt et al., 2024a). We used this demographic dataset to estimate average observed heterozygosity (H_O_) and its variance by sex, colony, sampling year and YOB, using a binary colony designation for colonies considered to be geographically separate based on infrequent observations of dispersal or migration (Chmura et al., 2016; Blumstein, 2025). Individual marmots and colonies were assigned to up-valley (n=4; Boulder, Marmot Meadow, North Picnic, Picnic) or down-valley colony (n=5; Avalanche, Bench, Gothic Town, Horse Mound, River) designations. We calculated each samples’ identity by descent probabilities (IBDp) and plotted IBDp by multidimensional scaling analysis mapping these colony designations in *PLINK* (MDS; --genome, --cluster and --MDS-plot 4). We further analyzed genetic structure by sex, colony and animal’s YOB by principal component analysis (PCA) in *FlashPCA* v2.1 (Abraham et al., 2017). We computed pairwise genetic differences between colonies and sexes as Weir and Cockerham’s *F*_ST_ in *VCFtools* v0.1.16, for which we compared weighted mean *F*_ST_ for the autosomes and sex chromosomes combined (vonHoldt et al., 2024a).

We analyzed genetic structure in *ADMIXTURE* using an unsupervised approach with maximum likelihood (Alexander et al., 2009). First, we assessed the likelihood (*Q*) of nine ancestral genetic partitions (*K*=2-10) based on their cross-validation error and considered the lowest error to represent the best fit. To resolve spatial patterns underlying these ancestry partitions, centroids were extracted from custom polygons designed over the area of each marmot study colony (Bobb & Adler et al., 2025) in the R package *sf*. We computed pairwise geographic distances (in km) and genetic distances (in F_ST_) between colonies before modeling a geographic cline to estimate transitions in ancestry along a user-defined, one-dimensional linear transect (Jofre & Rosenthal, 2021). We extended our transect 5 km north of the North Picnic colony and 5 km south of the River colony to span the full study region (Heppenheimer et al., 2018). Each sample was then positioned along the transect by projecting its centroid onto the transect and measuring Euclidean distances from the transect start point to each centroid’s projected point (Szymura & Barton, 1991). We used the transect to fit a sigmoidal cline with each sample’s *Q* value using nonlinear least squares regression in R. We extracted two mathematical parameters from the modeled cline to identify a putative genetic contact zone: the cline’s center or inflection point, the transect position where ancestry proportions changed most rapidly, and cline width, defined as pMax−pMin (*Q* values at the cline ends, estimated as free parameters instead of fixed tails) and divided by the maximum slope, the distance over which ancestry transitions were most likely to occur (Derryberry et al., 2014; Marshall et al., 2026).

Finally, we used *TreeMix* v.1.13 (Pickrell & Pritchard, 2012) to reconstruct maximum likelihood of plausible historical relationships between the study colonies while modeling up to 10 migration events (*m*=0-10), inferred from allele frequency covariance among colonies. We selected the number of migration events that maximized information gained across each incremental model, noting that the algorithm monotonically is a non-decreasing estimate. The marginal gain for each additional permitted migration event was used to determine the variance explained by the *m*^th^ edge (Evanno et al., 2005).

### Demographic history

The annual census of this marmot population since 1962 documented a significant increase in total number of individuals since at least the summer of 2000 (Ozgul et al., 2010; Blumstein, 2013; Blumstein, 2025). To model genomic signatures of this demographic change, we used the linkage-disequilibrium method in *GONE* (Santiago et al., 2020) to estimate effective population size (Ne) over the past 200 generations, with a focus on the most recent 50 generations. This method accounts for genetic drift and uses inferred recombination. We followed standard software recommendations and used the default parameters to disregard SNP pairs with a recombination rate >0.05, included loci on the first 200 scaffolds and assumed a constant genomic rate of recombination (1 cM per Mb). For Ne estimates, we used the minimally filtered SNP dataset and were conservative when interpreting “present-day Ne” as the most recent four generations (Novo et al., 2023). We converted the timescale from generations to calendar years based on 4-5 years per generation (Armitage, 2014), with a preference for 4 years per generation.

### Pedigree reconstruction and relatedness

For pedigree reconstruction and relatedness, we constructed a final, strictly filtered “pedigree” dataset from our demographic dataset, for which we further excluded loci with a minor allele frequency (MAF) <0.35 and >5% missingness per locus (--geno 5) in *PLINK*, requiring a 95% genotyping rate. We reformatted the pedigree dataset using --recodeA in *PLINK* to generate an additive genotype matrix, which we used as input to construct a high confidence pedigree based on parentage assignments in the R package *sequoia*. We imputed each sample’s sex and YOB for pedigree construction when known. We set a parentage assignment error rate of 0.5% and pedigree reconstruction (trio) error rate of 0.01% (Ritland, 1996; Lynch & Ritland, 1999). We visualized parentage assignments using the R package *kinship2* (Sinnwell et al., 2014) and pairwise pedigree relatedness using the *sequoia* function *GetRelM*.

### Inbreeding depression

We investigated the effect of genomic inbreeding on seasonal mass gain in pups, a life history trait that strongly predicts annual survival (Armitage, 2017; Paniw et al., 2020). To minimize effects of annual environmental variation, mass measurements were used for each animal in their first year of life (i.e., first year of data for individuals of known YOB, representing their body mass as a pup). Marmots with unknown YOB or insufficient mass measurements were excluded. From body mass measurements taken over the active season, we calculated an animal’s mass at two periods: their pup emergence mass, and 15 August mass (as a proxy for end-of-season mass). For 15 August mass, we used best linear unbiased predictors (BLUPs) to estimate mass on a given date (Jebb et al., 2021; Philson et al., 2022). Seasonal mass gain over an individual’s first year of life was calculated as 15 August mass minus emergence mass.

We then varied the genomic inbreeding predictor used to assess two metrics of inbreeding depression on seasonal mass gain: 1) an individual’s FROH, using their own inbreeding coefficient, and 2) combined parental FROH, using each parent’s inbreeding coefficients from known parent-offspring pairs in our pedigree. We fit linear mixed models (LMM) and tested the significance of each fixed effect using Satterthwaite’s approximation method in the R packages *lme4* (Bates et al., 2015), *car* (Fox & Weisberg, 2019), *tidyverse* (Wickham et al., 2019), *lmerTest* (Kuznetsova et al., 2017) and *performance* (Lüdecke et al., 2021). To test if an individual’s FROH predicted their seasonal mass gain as a pup, we fit the LMM: delta mass ∼ FROH + Sex + (1|Colony) + (1|YOB) for n=202 marmots across nine study colonies and 21 total YOBs. To test if the fixed effect of parental FROH predicted seasonal mass gain in their genetically confirmed offspring as a pup, we fit the LMM: delta mass ∼ FROH_dam_ + FROH_sire_ + Sex + (1|Colony) + (1|YOB), for n=44 marmots across eight study colonies and four total YOBs. We included YOB as a random effect to represent generational cohorts and as a centered number, and animal’s colony as a random effect to account for non-independence among individuals from the same colony and YOB. We assessed collinearity among each parent’s genomic FROH using the variance inflation factor (VIF).

## Results

We obtained genome-wide genotype data for 284 marmots. Prior to processing and filtering, we obtained 808,165 loci at a mean sequencing depth of 48.6-fold coverage (s.d.=51.9) and an average nucleotide diversity of *π*=0.192. After filtering, we retained 705,360 loci with at least 10-fold sequencing coverage, of which we retained 153,550 loci within the minimally filtered dataset (total genotyping rate=94.4%) and excluded 15 individuals due to high (>20%) SNP missingness. We constructed our filtered demographic dataset from 269 marmots genotyped at 17,393 statistically unlinked and neutral loci, after removing another 135,474 loci that exceeded the genotypic Pearson’s correlation threshold and 683 loci which deviated from Hardy-Weinberg Equilibrium (total genotyping rate=95.5%). We constructed our pedigree dataset from 1,902 loci from the same number of individuals after removing another 7,755 loci which exceeded the genotype missingness threshold of 5% and 7,736 loci with a high minor allele frequency (>0.35). We found moderately high H_O_ levels across SNPs in our demographic dataset (mean ± s.d.=0.28 ± 0.03; Table S1) and evidence of population structure by geographic isolation (Figure 1A).

**Figure 1.**
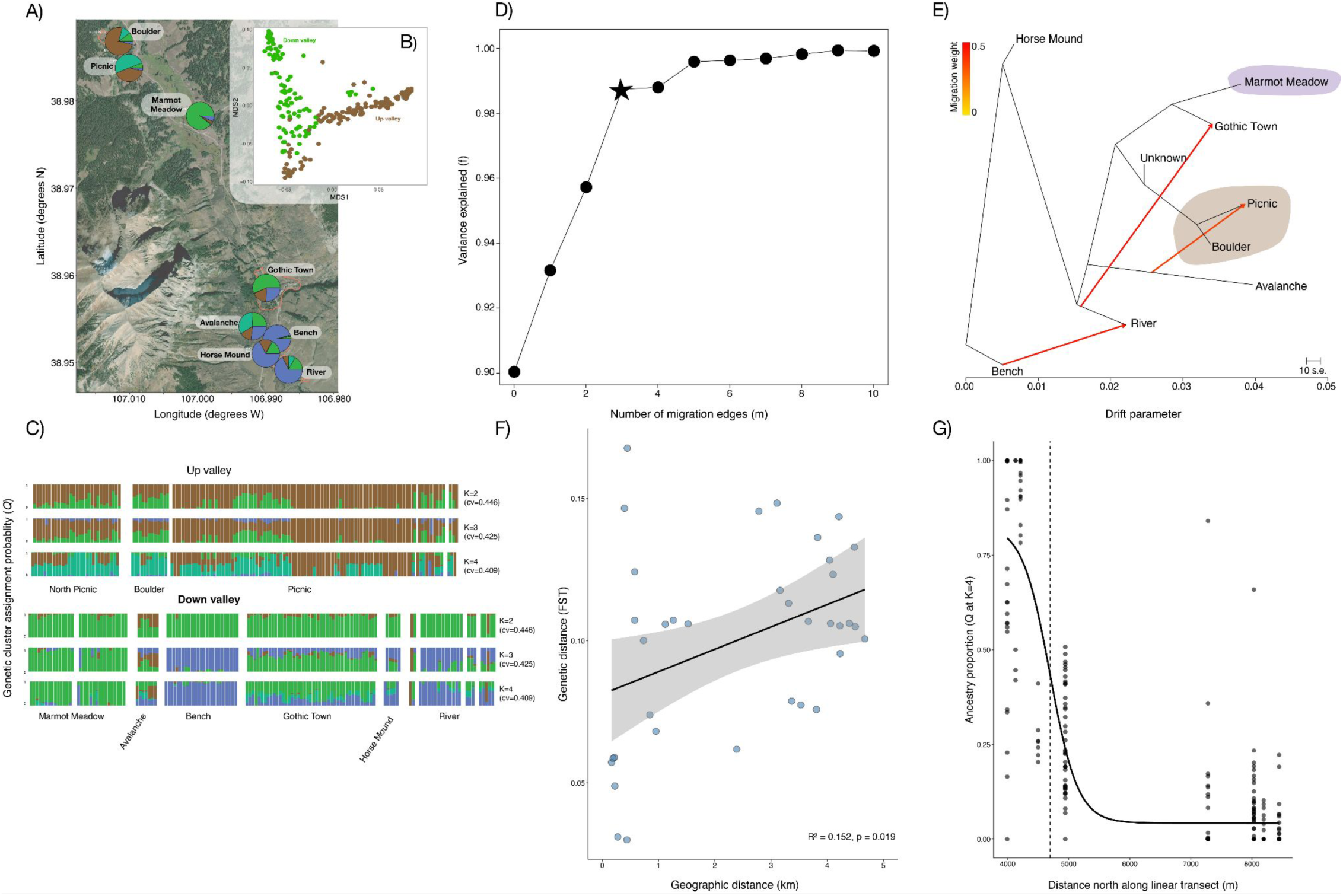
**>A)** Map of the study area. Pie charts show mean proportions of population membership from unsupervised admixture at k=4 genetic groups; individual colony names are provided. **B)** Multidimensional scaling analysis of demographic filtered SNPs for 286 marmots, indicating each sample’s up- or down-valley colony designation. **C)** Genetic cluster assignment probabilities (*Q*) to each genetic partition (*K*=2-4) with colony names and estimated cross-validation (cv) error. **D)** Variance explained for each modeled migration event and the selected number of migration events (starred), after which marginal gain in explained variance plateaus. **E)** Relationships between geographic colonies and three putative migration events with directional arrows. Color bar indicates migration edge weight, and branch lengths indicate migration strength proportional to drift. **F)** Genetic distance by geographic isolation for individual colonies from pairwise mean F_ST_. **G)** Logistic geographic cline for ancestry proportions (*Q* at K=4) modeled along a linear transect spanning the study area; dashed line indicates the cline’s center (inflection point).

### Genetic isolation is enforced by colony geography with limited south-to-north connectivity

Our northernmost study colony, North Picnic, was located 4.669 km from the southernmost colony, River, while the two nearest colonies, Horse Mound and Bench, were separated by only 215.339 m. Multidimensional scaling analysis of genetic structure showed initial groupings by valley position alone (Figure 1B). Average observed mean heterozygosity (H_O_) also differed by valley position (down- valley=0.283; up-valley=0.278 (1-tailed *t*-test; *p*=0.0803, Wilcoxon rank sum test; W=10,588, *p*=0.001). We observed no genetic structure in PCA by sex, and YOB was significant for PC2 only (*F*=1.797, *p*=0.018), but individual colony assignments were highly significant for PC1 and PC2 (PC1: *F*=105.4, *p*<2×10^-16^; PC2: *F*=102.6, *p*<2×10^-16^) (Figure S1). Using an unsupervised approach with diminishing cross-validation errors, we found optimal ancestry values for *Q* at K=4 (Figure 1C; Table S2). With fewer genetic partitions, ancestry structure approximated the a priori up- and down-valley valley designations, but with increasing number of partitions, a tri-part substructure emerged (Figure S2), supported by the individual colony PCA clustering (e.g., Marmot Meadow was distinct from up-valley Picnic and Boulder and more similar to the down-valley colony Gothic Town).

Using a genetic migration detection tree reconstruction model, we found the best support for three migration events (Figure 1D). This bifurcating topology emerged from the earliest split into the two down-valley colony lineages (Figure 1E). Gothic Town represented a transition of both genetic partitions and was more similar in ancestry to two of the up-valley colonies. The longer branches of nearly all colonies relative to Bench and Horse Mound indicate their potentially smaller long-term Ne or geographic isolation. Our inferred three migration events have strong weights depicting unidirectional migration directly between adjacent colonies (e.g., Bench to River), while the other two migration events have longer edges, implying gene flow between geographically distant colonies and restricted from north to south (Figure 1E). Genome-wide weighted *F*_ST_ values supported significant genetic differentiation between individual colonies by distance (linear regression, F(1,34)=6.096, multiple R^2^=0.152; *p*=0.019) (Figure 1F; Table S3). We found less genetic differentiation solely by colony designation (weighted estimate, *F*_ST_=0.047) and minimal differentiation by sex (weighted estimate, *F*_ST_=0.002). Average shared ancestry by sample (*Q* at cluster 4, K=4; at K=2 the sigmoidal model failed to converge) declined sharply from south to north along our modeled geographic cline (Figure 1G). The cline’s inflection point was at 5,648.579 m along the linear transect roughly parallel to County Road 317 and had a width of 914.03 m. Both the Gothic Town and Avalanche colonies fell within the modeled cline’s contact zone (Figure S3). Colonies south of Avalanche and Gothic Town were dominated by shared ancestry, while sites north of the cline’s inflection point had low (Q=∼0.04) shared ancestry (Figure 1G).

### A recent reduction in census and effective population size aligns with temporal inbreeding trends

Trapping data revealed a decline to ∼100 individuals in 2011, followed by a gradual increase to ∼200 as of summer 2025 (Figure 2A). The Picnic colony had the highest long-term census size (n=99), while the Horse Mound colony had the lowest (n=6; Table S1). Using 131,850 out of 153,550 loci (85.9%) genotyped at high confidence, we found that Ne rose gradually from 50 to 17 generations ago (Ne=9840 and 16793.8, respectively), followed by a steep recent decline, approaching a present-day estimate of Ne=53.2 (generations 1-4). Translating generation time into calendar years, Ne decline began between 1997 (generation 8, Ne=13380.5) and 2009 (generation 5 Ne=356.7) (Figure 2B). We identified variable levels of genomic inbreeding from autozygosity, with most individuals showing low inbreeding and a long right tail (mean ± s.d.=0.11 ± 0.08, range=0.02-0.45) (Figure 2C). We identified 50,173 ROH blocks and an average 185.1 ROH blocks per animal (s.d.=43.3). Using two individual-based metrics of genetic diversity over time, H_O_ and FROH, we found that H_O_ had no temporal trend (Pearson’s product-moment correlation, R=−0.205, p=0.296) and that FROH increased by YOB when all samples were included (R=0.18, p=0.004), but not when pups born in 2025 were removed (R=0.049, p=0.505). In living marmots sampled in each year of the study, annual mean H_O_ reached minimum values post-2010, coinciding with high annual mean FROH, before reaching their present-day minimum and maximum annual means, respectively (2025: H_O_=0.266; mean FROH=0.144), driven by pup sampling (Figure 2D; Figure 2E). Males on average were more inbred than females (Wilcoxon rank sum test, continuity correction; mean FROH males=0.134, females=0.086; W=13454, *p*=3.24×10^-13^). There was no difference in inbreeding by colony designation (mean FROH up-valley=0.106, down-valley=0.107; W=7790.5, *p*=0.222).

**Figure 2.**
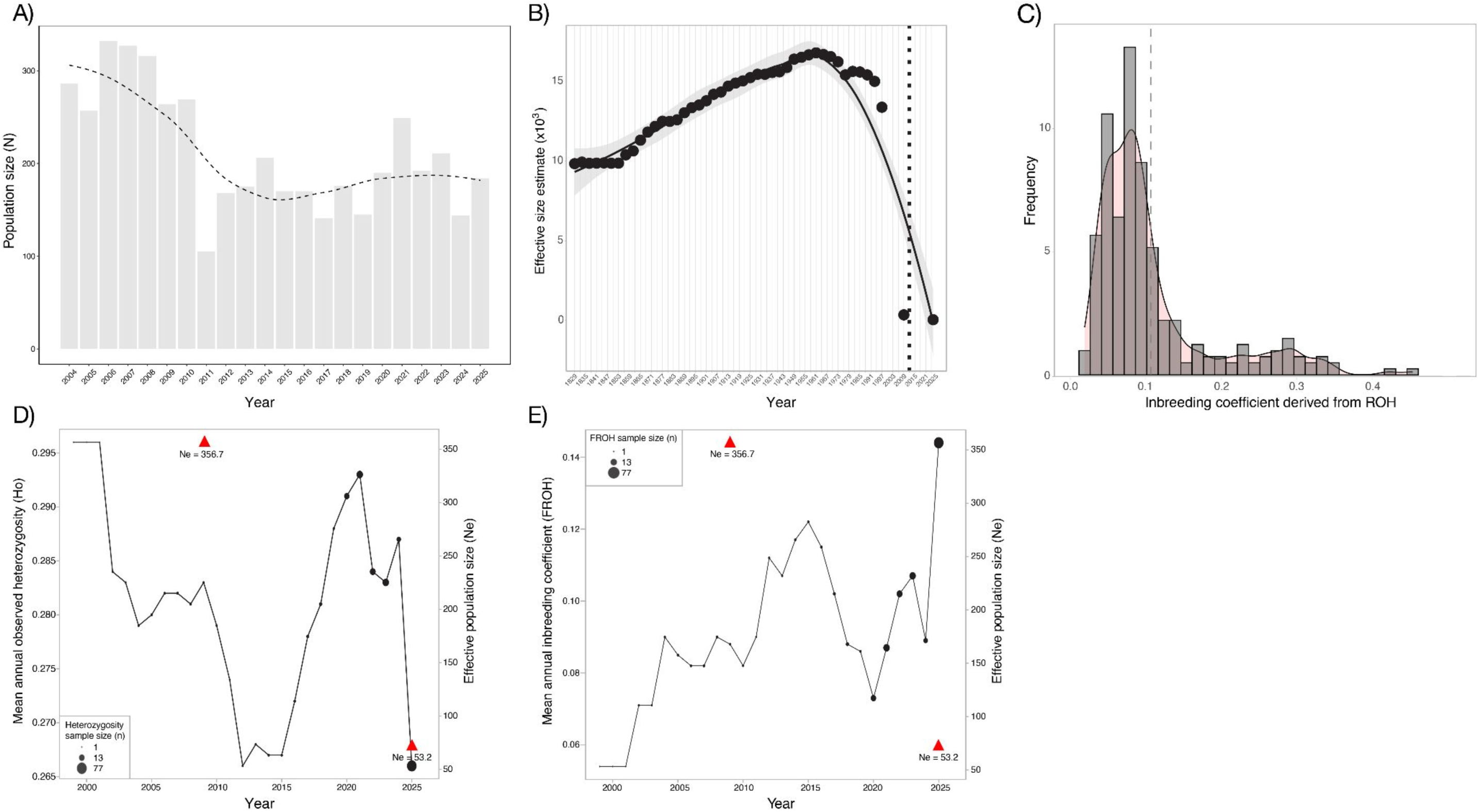
**A)** Census size over the study period (2004-2025). **B)** Ne estimated from linkage disequilibrium (4 years per generation); vertical dashed line indicates year of observed census decline from (A), 2011. **C)** Genomic inbreeding values derived from autozygosity (ROH) (dashed line indicates mean ROH with the smoothed density function shaded in pink). **D)** Annual mean observed heterozygosity (H_O_), and **E)** Inbreeding coefficient (FROH) in living animals sampled each year over the study period, where red triangles and the y-axis show the Ne estimates in 2009 and 2025, respectively, the two generations that overlap with genetic sampling. Point size indicates the number of animals sampled in each calendar year.

### Pedigree reconstruction indicates minimal dispersal, reinforcing geographic isolation

We reconstructed 134 dyads and 112 trios for 271 marmots using genotype data at 1,902 SNPs, connected in the pedigree by 29 female and 37 male placeholders for an additional 10 dyads and 16 trios (Figure 3). We identified 44 dams and 10 sires, which directly accounted for 240 pups directly connected in the pedigree, not including placeholders or founders themselves (Table S1). Based on this pedigree, our pairwise relatedness matrix defined eight detailed genealogical relationships (n: self=272, maternal=260, paternal=259, full sibling=378, half sibling=1096, grandparent=487, and avunculate=1096) (Figure S4). After excluding self-pairs, we found the average pedigree-based relatedness values within a colony were significantly higher than between colonies (0.11 and 0.01, respectively; Welch’s two-sample t-test: t=-85.2, df=20728, *p*<2.2×10^-16^) (Table S4). We also identified variation in relatedness by colony. Based on the pedigree, Picnic (up valley, r=0.10) and Gothic Town (down valley, r=0.08) had the lowest within-colony average relatedness, while Avalanche (down valley, r=0.423) had the highest relatedness (Table S4). We analyzed two focal colonies: the down-valley Bench colony (Figure 4A) and up-valley Picnic colony (Figure 4B) as example of reconstructed multi-generational pedigrees and found that these two colonies depicted several events of colony fidelity, dispersal or inter-colony mating. However, no dispersal or mating events occurred directly between these two example geographic colonies.

**Figure 3.**
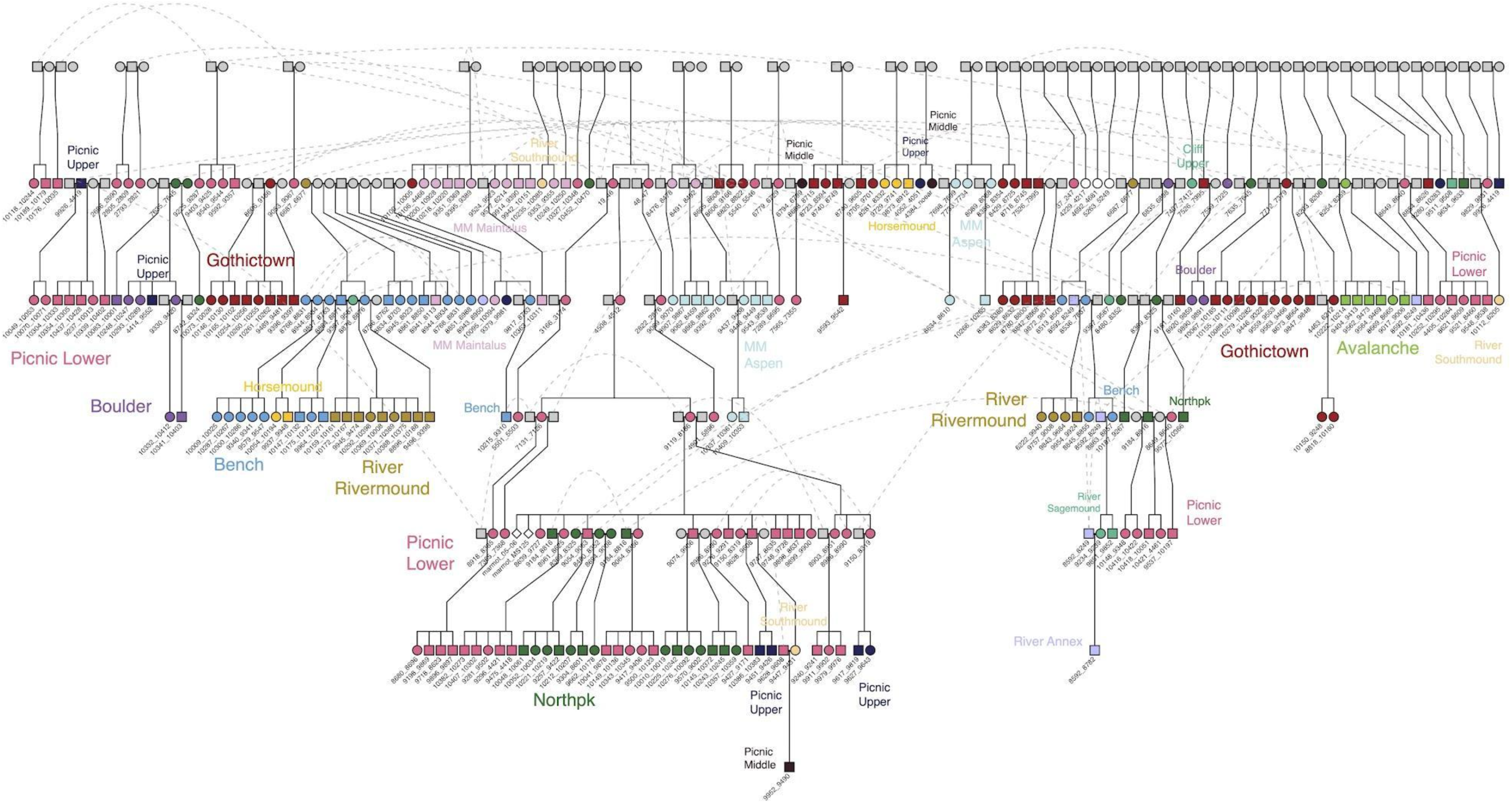
Reconstructed genetic pedigree from 1,902 pedigree filtered SNPs. Each animal’s unique identifying ID is provided in the pedigree. Dashed lines indicate a marmot is found in different locations of the pedigree space with respect to genetically inferred reproduction. Colored symbols indicate geographic colony identity with name. (Symbols: females, circles; males, squares). (Abbreviations: MM, Marmot Meadow).

**Figure 4.**
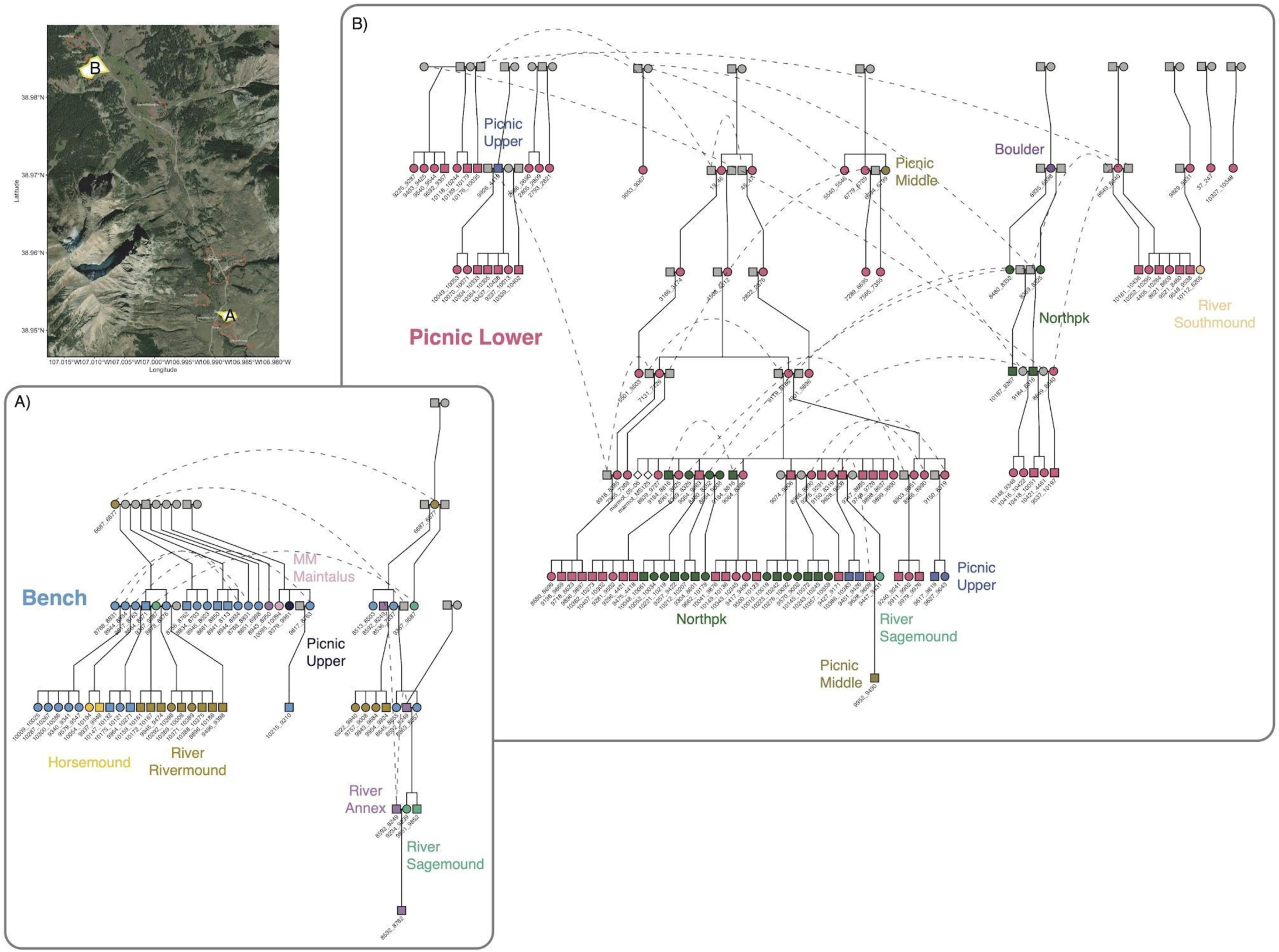
Examples of a multigenerational, colony-specific pedigree for the **A)** down-valley Bench colony, and **B)** the up-valley Picnic colony, geographically separated by 4.04 km in the East River Valley. Dashed lines indicate when an individual is sampled in a different geographic colony of the pedigree space with respect to genetically inferred reproduction. Colored symbols indicate colony and burrow membership. (Symbols: females, circles; males, squares). (Abbreviations: MM, Marmot Meadow).

### Parental inbreeding is associated with seasonal mass gain in offspring

Genomic breeding was associated with seasonal mass gain in an individual’s first year of life when colony and YOB were not modeled (ANOVA; FROH values were log-transformed to correct for non-normality of autozygosity values, F₁,₂₀₀=5.78, *p*=0.017; Figure S5; Figure S6). To determine if this relationship was significant after accounting for an individual’s colony and YOB, we fit an LMM with FROH and sex as fixed effects and their colony and YOB as random intercepts. In this model, FROH did not predict mass gain (β=−70.7, SE=179.0, t₁₈₇.₇=−0.40, p=0.693), but males gained more mass than females (β=92.3 g, SE=27.2, t₁₇₉.₄=3.40, *p*<0.001). Random-effect standard deviations were similar for colony (169.8 g), YOB (135.3 g), and residual individual-level variation (162.3 g). To test if parental inbreeding predicted seasonal mass gain in their genetically confirmed offspring, we fit an LMM with maternal FROH, paternal FROH and offspring sex as fixed effects, and with colony and YOB as random intercepts. Maternal and paternal FROH were not collinear based on VIF for resolved parent-offspring pairs. Maternal FROH was negatively associated with seasonal mass gain in their offspring (β =−10,914, SE=3,220, t₃₁.₅=−3.39, *p*=0.002), while paternal FROH was positively associated with mass gain (β=14,222, SE=6,212, t₂₅.₂=2.29, *p*=0.031; Figure S6). Offspring sex had no effect on mass gain in this model (β=65.4 g, SE=41.2, t₂₇.₃=1.59, *p*=0.124), but colony was significant and accounted for more variation (s.d.=358.0 g) than YOB (s.d.=162.5 g) or residual individual-level variation (s.d.=112.7 g).

## Discussion

Using longitudinal data from a wild marmot population over more than two decades, we quantified genetic diversity, relatedness and inbreeding to examine deviations from historical trends and their fitness implications. Alongside evidence of genetic structure by fine-scale geographic isolation and discovery of a recent and lasting population decline, we found genomic metrics of inbreeding have increased and were associated with a reduction in seasonal mass gain in the offspring of inbred parents. While these marmot colonies have previously seen high rates of gene flow (Schwartz & Armitage, 1980) and population expansion (Ozgul et al., 2010), this recent precipitous decline suggests these changes to demography may have produced new consequences to genetic diversity, and potentially to reproductive fitness.

### Genetic structure across long-term colonies reveals a latitudinal pattern to gene flow

Our genomic data reflect a pattern of isolation by distance with cryptic barriers to gene flow enforcing population subdivision. While dispersal and migration are common at highly proximal scales, increased geographic distance and heterogenous terrain create genetic differentiation on the landscape. Across the East River Valley, this marmot population formed three genetic groups supported by MDS, PCA and *F*_ST_, and which deviated from previous geographic distinctions of “up-valley” versus “down-valley”. Most notably, “down-valley” Gothic Town which hosts the RMBL station, is more genetically similar to the Marmot Meadow colony, previously assumed to be an isolated “up-valley” colony from observations and trapping data (Svendsen 1974; Armitage, 1991). This finding supports our study hypothesis that Gothic Town is a critical location for gene flow between otherwise isolated colonies; indeed, only one predicted migration event was recorded between any “up-valley” and “down-valley” colonies (from Avalanche to Picnic; Figure 1E). This hypothesis was further supported by a modeled geographic cline, which indicated Gothic Town is within a narrow contact zone along a south to north gradient in ancestry. Unidirectional gene flow indicates marmot dispersal may align with climate stressors, as southern colonies have become warmer and more likely to be snow-free in recent decades (Cordes et al., 2020). This restricted pattern of gene flow from south to north suggests that recent generations may be dispersing exclusively to northern, higher elevation sites, a trend observed in species distribution modeling in other subalpine species subject to climate-induced range shifts with warming (Mathewson et al., 2017; Simma et al., 2025).

Ultimately, we found minimal gene flow between distal colonies with rare long-distance dispersal, shedding light on sex-biased migration events, which are challenging to observe directly. Terrain features such as waterways, roads or human presence may inhibit longer-distance dispersal in marmots (Faulkner, Hulme & Wilson, 2024; Marshall et al., 2026), potentially increasing inbreeding at isolated sites. These long-term study colonies also varied in plant abundance, hibernaculum availability (Duporge et al., 2026) and insulation from predators, influencing annual census sizes (Blumstein et al., 2007) and connectivity. Integrating genetic information suggests that small home ranges and limited dispersal based on these factors could be more important in shaping fitness outcomes in marmots than presumed differences in environmental harshness by elevation, given the southernmost colony, River, at 2,867 m, is only 125 m lower than the northernmost colony, Picnic (2,992 m; Oli & Armitage, 2004).

Subtle genetic structure is consistent with the social and mating structure of this marmot species. Our ∼5 generation pedigree and relatedness estimates showed mating does occur between distant colonies, despite being rarely documented (Edic, 2020), yet ancestry structure is maintained (Figure 1A). In marmots, social hierarchy is reinforced by matrilineal dominance, in which female yearlings remain philopatric and yearling males tend to emigrate (Downhower & Armitage, 1981; Lenihan & Van Vuren, 1996; Schwartz, Armitage, & Van Vuren, 1998; Armitage et al., 2011). Adult reproductive females thus remain near natal burrows and reproduce with close kin (Armitage, 1991; Schwartz & Armitage, 1980; Blumstein, 2025), generating persistent patterns of reduced heterozygosity (Olson et al., 2012), which we also observed in our genomic inbreeding data. Yet female philopatry is primarily related to the social environment, such as continued local presence of the dam and frequency of non-aggressive interactions with matrilineal kin, rather than by inbreeding avoidance or direct competition (Armitage et al., 2011). Within colonies where long-standing matrilines have been disrupted by major overwinter die-offs, demographic stochasticity could have altered the social hierarchy enforcing female philopatry.

### Increased inbreeding and a slow recovery of genetic diversity since recent demographic collapse

Marmots have been present at high (>2,800 m) elevations in the Upper Gunnison Basin for at least the last 49,000 years (Reynard et al., 2015; Emslie & Meltzer, 2019; Frey, 2026). We found that this local marmot population experienced consistent growth in Ne since the 19th century. Paired census and genomic data indicate the population underwent its decline beginning prior to the largest overwinter die-off recorded in over 60 years of annual study following the winter of 2010. This demographic change (from ∼300 to ∼200 individuals) was accompanied by increased inbreeding and decreased heterozygosity in contemporary generations. Such a demographic shift may signal the inability for this small and isolated population to rebound from a disruption to a previously stable Ne. Our results provide the first evidence of a potential demographic collapse stemming from this die-off event, possibly exacerbated by reduced adult survival in adverse environmental conditions with climate change (Paniw et al., 2020). The winter of 2025, one of the driest, warmest water years on record in western North America, coincided with the second largest overwinter die-off in study history (unpublished data), suggesting that historical changes in winter snowpack could predict future census size, but likely with limits to predictability.

### Intergenerational effects of genomic inbreeding on a trait predictive of annual survival

Inbreeding can persist despite fitness losses when cost of forgoing reproduction is high (Waser et al., 1986; Kokko & Ots, 2006), particularly when environmental conditions are unpredictable (Petrullo et al., 2025). Few studies have investigated relationships between inbreeding and early-life growth despite their critical role in fitness (Young, 1976; Carrier et al., 2022; Wells et al., 2020; Allison et al., 2026). In marmots, we found that higher levels of maternal inbreeding reduced offspring mass gain in their first year of life. Our results contrast one similar study in Antarctic fur seals (*Arctocephalus gazella*), which found no relationship between individual or maternal inbreeding on pup birth mass (Paijmans et al., 2024) but align with theoretical expectations for inbreeding depression in offspring using ROH (Kyriazis, Robinson, & Lohmueller, 2025). In further alignment with our results, previous work on inbreeding in this marmot population also found a fitness-related effect at the offspring rather than the individual level (Olson et al., 2012). Although our seasonal mass gain and inbreeding models differed in direction and magnitude, these results suggest that inbreeding can influence early-life growth and may indicate inbreeding depression in marmots.

Given our modest model sizes, we interpret these results with caution. We analyzed an unbalanced sample set, with fewer paternal than maternal parentage assignments (43 dams; 10 sires). We also found that individual inbreeding had no effect on seasonal mass gain and that pup colony explained more variance in mass gain than their birth year. Our results together support stronger effects of birth date, weaning date, maternal provisioning and environmental contingency on seasonal mass gain, which may each play a larger role on pup fitness than genomic inbreeding. Maternal provisioning is strongly correlated with seasonal mass gain in marmot offspring (Ozgul et al., 2010) due to annual plasticity in birth and weaning date, which are strongly influenced by annual timing of reproduction to environmental conditions (St. Lawrence et al., 2023). If dams can reproduce and wean pups earlier, their offspring can achieve a higher seasonal body mass by the end of the season (Ozgul et al., 2010; St. Lawrence et al., 2022). In support of an outsized role for maternal provisioning, older females produce daughters with higher lifetime fitness, irrespective of annual fluctuations in environment quality by birth year (Kroeger et al., 2020). Selection against inbreeding may also be enacted later in life, beyond pup mass.

## Conclusions

Our results highlight the utility of integrating genomic datasets with traditional population monitoring to understand the multifaceted mechanisms underlying demographic change. We found this marmot population to be in a moment of sudden decline, suggesting an uncertain future with climate change. The role of the Gothic Town colony, the site of the RMBL, in facilitating gene flow between otherwise isolated colonies is of particular relevance with respect to environmental change. With ever-increasing tourism and annual human presence at the RMBL, the behavior of marmots in this more “urbanized” colony site has been altered (Adler et al., 2026), likely alongside other traits and resident species. Wildlife in cities have been found to possess genetic and phenotypic changes specific to urban settings (Alberti et al., 2017; Schell et al., 2021), but the effects of human activity on wildlife outside highly urbanized ecosystems are still poorly understood (Tranquillo et al., 2024). Our study takes a deep dive into one study species and population, but our findings on cryptic barriers to dispersal, Ne dynamics and potential for inbreeding depression offer relevant takeaways to other natural populations of Least Concern, many of which are currently in decline. We expand on a wealth of existing knowledge on this marmot system, identifying long-term drivers of genetic isolation and documenting recent reductions in population size that can be used to understand and predict future risk of quasi-extinction or extirpation.

## Author Contributions

Conceptualization: SRT, BVH; Formal Analysis: all authors; Visualization: BVH, SRT; Data Curation: all authors; Resources: BVH; Supervision: SRT, BVH; Funding Acquisition: BVH; DTB; JGAM, Writing, original draft: SRT; Writing, review and editing: all authors.

## Acknowledgements

We are grateful to the many decades of marmoteers who collected these data in the field and to the staff at the Rocky Mountain Biological Laboratory for logistical support with data collection. We thank Dr. Craig Marshall and Augustin Birot for helpful discussions on manuscript analyses. We thank Dr. Yeraldi Loera for sharing computer code and members of the vonHoldt lab for feedback that improved this manuscript.

## Ethics Statement

Wild yellow-bellied marmots from the East River Valley, Gothic, Colorado, USA were captured and handled under the following permit: Colorado Parks and Wildlife (Permit Number: TR917). All animal related procedures were ethically reviewed and approved annually by IACUC at the University of California Los Angeles (protocol 2001-191-01) and the Rocky Mountain Biological Laboratory (RMBL).

## Data Availability Statement

Trimmed and aligned sequencing data (accession numbers listed in Table S1) are deposited on NCBI Sequence Read Archive (SRA; https://www.ncbi.nlm.nih.gov/sra) under BioProject PRJNA1555184.

