## Supplemental Material for "Spatial and temporal drivers of genomic inbreeding in a declining population of yellow-bellied marmots"

**Supplementary Material** **for “***Spatial and temporal drivers of genomic inbreeding in a declining population of yellow-bellied marmots.***”**

**Supplemental Table S1.** Sample metadata with NCBI SRA accession numbers (see file TableS1.xlsx).

**Supplemental Table S2.** Cross-validation errors for each genetic partition (K) tested for ancestry values*.*

| Genetic partitions | Cross-validation error |
| --- | --- |
| K=2 | 0.446 |
| K=3 | 0.425 |
| K=4 | 0.409 |
| K=5 | 0.399 |
| K=6 | 0.390 |
| K=7 | 0.386 |
| K=8 | 0.377 |
| K=9 | 0.374 |
| K=10 | 0.371 |

**Supplemental Table S3.** Pairwise F_ST_ values across the nine long-term marmot study colonies.

| Colony 1 | Colony 2 | F_ST_ |
| --- | --- | --- |
| Picnic | North Picnic | 0.030 |
| Picnic | Boulder | 0.057 |
| Picnic | Marmot Meadow | 0.106 |
| Picnic | Gothic Town | 0.079 |
| Picnic | Bench | 0.128 |
| Picnic | Horse Mound | 0.106 |
| Picnic | Avalanche | 0.107 |
| Picnic | River | 0.095 |
| North Picnic | Boulder | 0.031 |
| North Picnic | Marmot Meadow | 0.106 |
| North Picnic | Gothic Town | 0.076 |
| North Picnic | Bench | 0.133 |
| North Picnic | Horse Mound | 0.105 |
| North Picnic | Avalanche | 0.123 |
| North Picnic | River | 0.101 |
| Boulder | Marmot Meadow | 0.107 |
| Boulder | Gothic Town | 0.077 |
| Boulder | Bench | 0.144 |
| Boulder | Horse Mound | 0.105 |
| Boulder | Avalanche | 0.136 |
| Boulder | River | 0.106 |
| Marmot Meadow | Gothic Town | 0.062 |
| Marmot Meadow | Bench | 0.148 |
| Marmot Meadow | Horse Mound | 0.118 |
| Marmot Meadow | Avalanche | 0.146 |
| Marmot Meadow | River | 0.113 |
| Gothic Town | Bench | 0.100 |
| Gothic Town | Horse Mound | 0.074 |
| Gothic Town | Avalanche | 0.107 |
| Gothic Town | River | 0.068 |
| Bench | Horse Mound | 0.059 |
| Bench | Avalanche | 0.168 |
| Bench | River | 0.049 |
| Horse Mound | Avalanche | 0.147 |
| Horse Mound | River | 0.059 |
| Avalanche | River | 0.124 |

### **Supplemental Table S4.** Descriptive statistics for pedigree-based individual pairwise relatedness within and between marmot colonies, as well as within-colony relatedness comparisons.

|  |  | Pedigree-based relatedness descriptives: | |
| --- | --- | --- | --- |
| Comparison | Number of colony pairs | Mean | Standard error |
| Between | 53,296 | 0.012 | 0.0003 |
| Within | 18,796 | 0.109 | 0.0011 |
| *Within colony* | | | |
| Avalanche | 42 | 0.423 | 0.0215 |
| Bench | 552 | 0.293 | 0.0076 |
| Horse Mound | 20 | 0.219 | 0.0626 |
| River | 552 | 0.201 | 0.0095 |
| Marmot Meadow | 930 | 0.177 | 0.0065 |
| Boulder | 132 | 0.110 | 0.0139 |
| Picnic | 14,762 | 0.097 | 0.0011 |
| Gothic Town | 1,806 | 0.080 | 0.0036 |

**Supplemental Figure S1.** Population genetic structure of 269 marmots genotyped at 17,393 statistically neutral and unlinked loci inferred from principal component analysis (PCA), by (A) sex (F: female or M: male), (B) year of birth, known for all except four marmots in the study period (range: 1999-2025), and (C) resident geographic colony from trapping records.

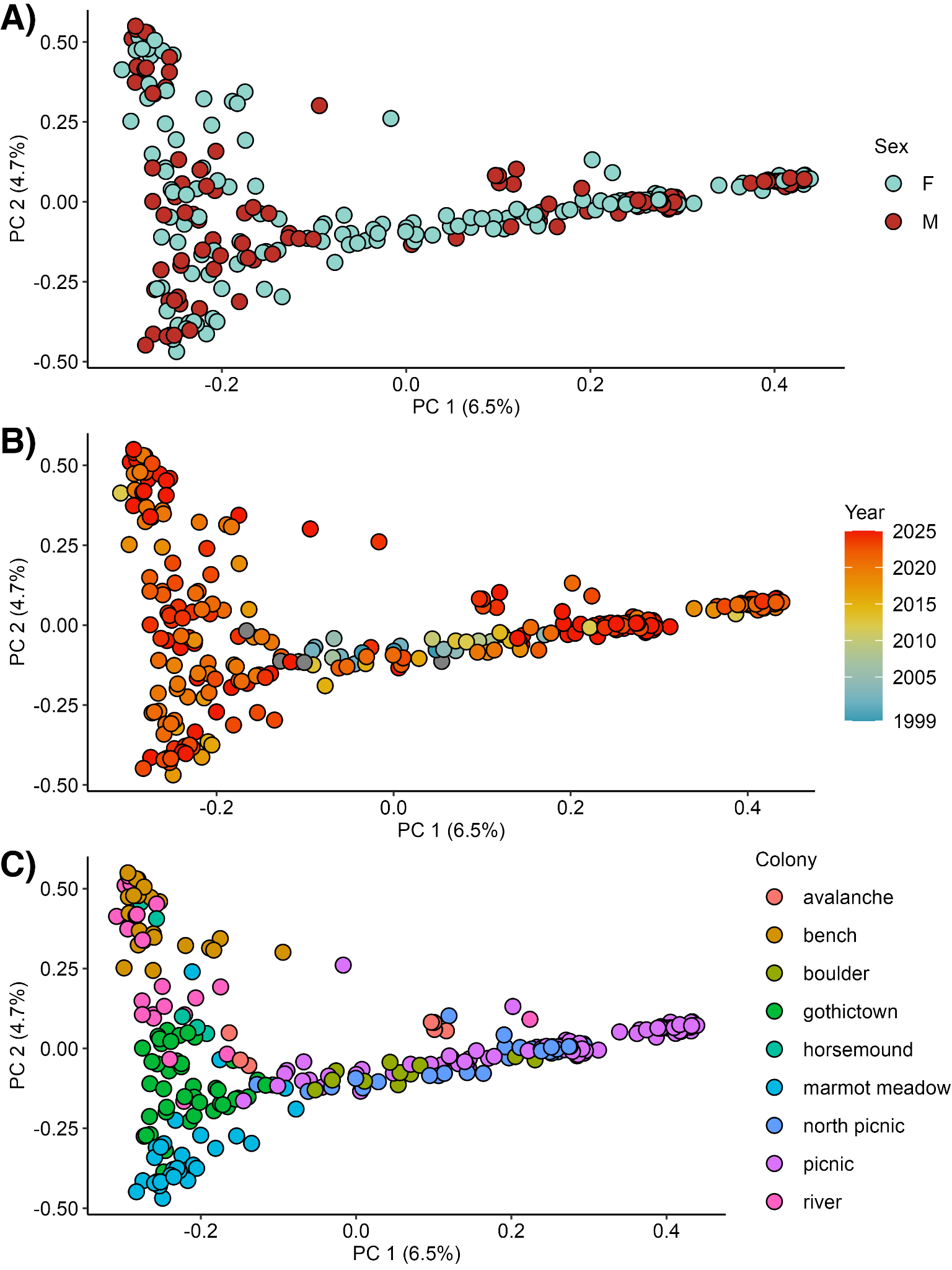

**Supplemental Figure S2.** Genetic cluster assignment probabilities (*Q*) showing each unsupervised admixture genetic partition (*K*=2-10) with colony name and cross-validation (cv) error estimate provided. Up- and down-valley colony designations are specified above individual colonies in each designated area.

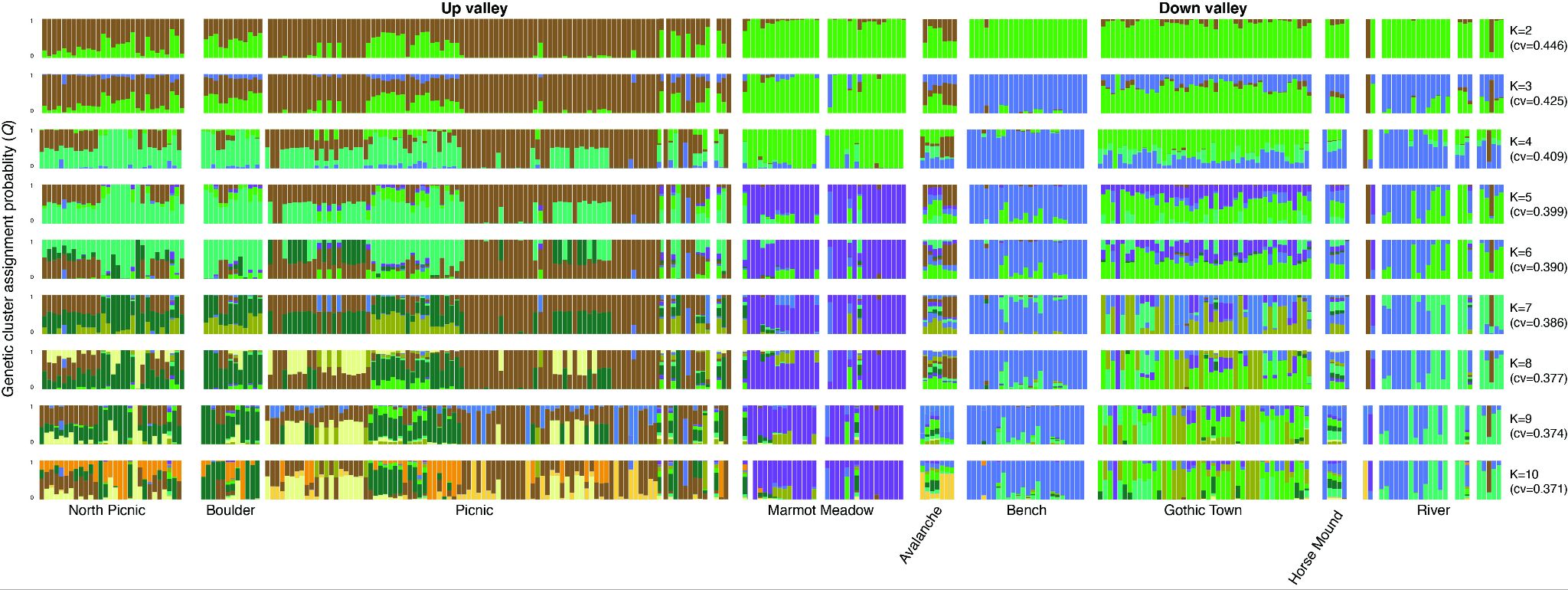

**Supplemental Figure S3.** Linear transect encompassing the study area used to model a geographic cline produced from centroid data from custom WGS84 polygons. Longitude spacing on the x-axis was increased for visualization. The northernmost colony centroid, North Picnic (NP) and southernmost colony centroid, River (R) were used to create the one-dimensional linear transect. All colony centroids are plotted in blue. Transect end points (plotted in red) were extended 5 km directly north and south of the NP and R centroids, respectively, to ensure no colony centroids overlapped or within the linear transect itself. The transect length itself was selected randomly and is arbitrary to the cline analysis. Using these end points, a line was generated as a *sf* “line” object. The orange boundary box (1000 m length was selected for ease of visualization, the length of the genetic cline was not modeled) was created using the output modeled cline’s center point along the transect (point of steepest ancestry inflection) and the cline’s width (914 m north and south of the center point, orange dashed lines) along the linear transect.

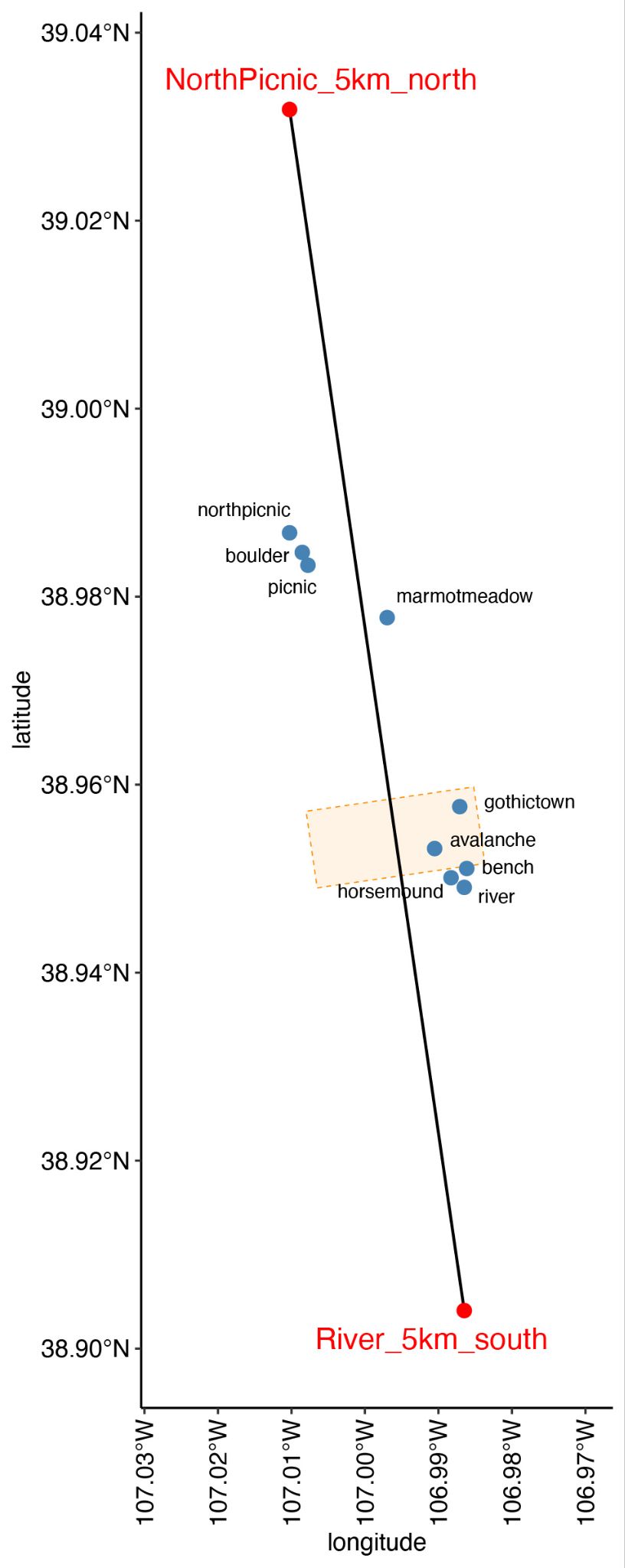

**Supplemental Figure S4.** Pairwise relatedness with relationship categories color coded from inferred pedigree structure (sample size per category provided in parentheses). Sample identities are listed on each axis, including any unknown parents (e.g., MXXXX placeholder sires; FXXXX, placeholder dams).

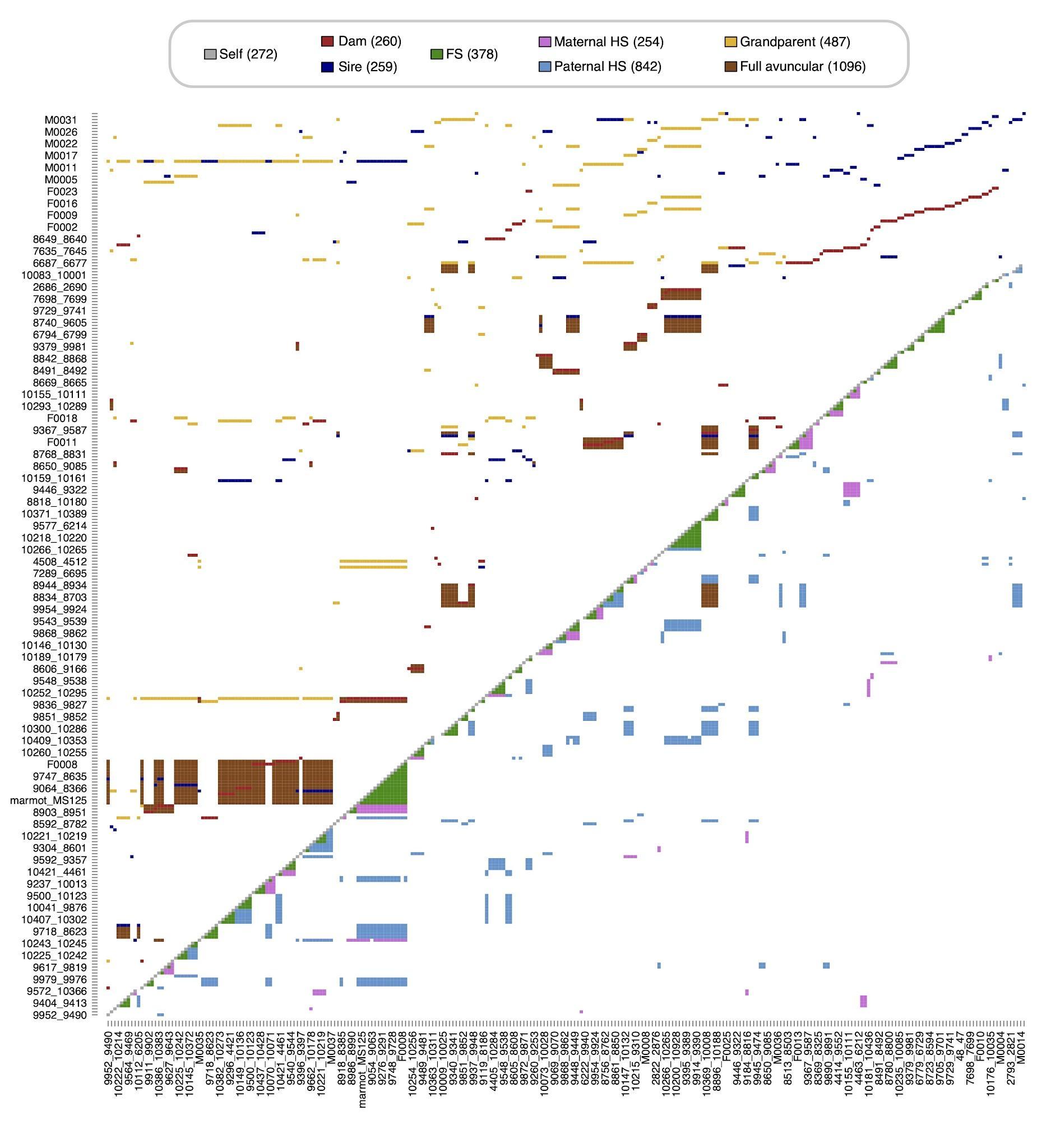

**Supplemental Figure S5.** ANOVA and quantile-quantile (QQ) plots before (top) and after (bottom) log transformation of genomic inbreeding coefficient (FROH) for linear mixed models. Left: histogram of residual value frequencies before and after transformation showing pre-transformed skew in negative values; Right: QQ plots of sample quantiles compared to a uniform distribution before and after transformation (95% CI is shaded), showing reduced inflation in log transformed values. Log transformation improved but did not entirely alleviate non-normality of FROH data (Shapiro-Wilk test, W=0.98517, *p*=0.03234).

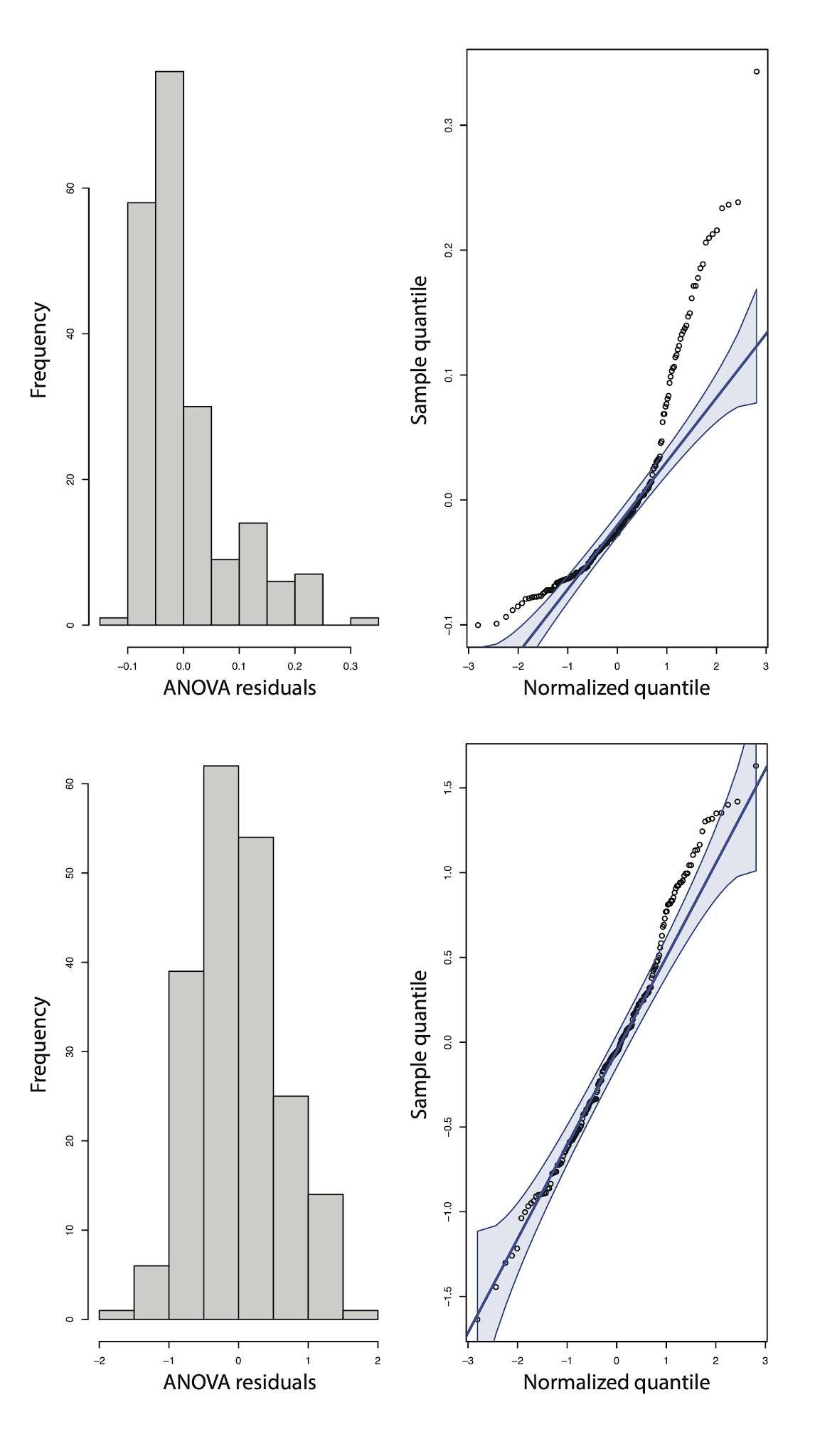

**Supplemental Figure S6.** **A)** Estimated seasonal mass gain (g) between emergence and 15 August in an individual’s first year of life as a function of individual genomic inbreeding coefficient (FROH). **B)** Offspring seasonal mass gain (g) between emergence and 15 August as a function of the genomic FROH of each parent, with genetically confirmed offspring from pedigree reconstruction. Linear regression correlation coefficients and p-values are shown; statistics in the main text are from linear mixed models.

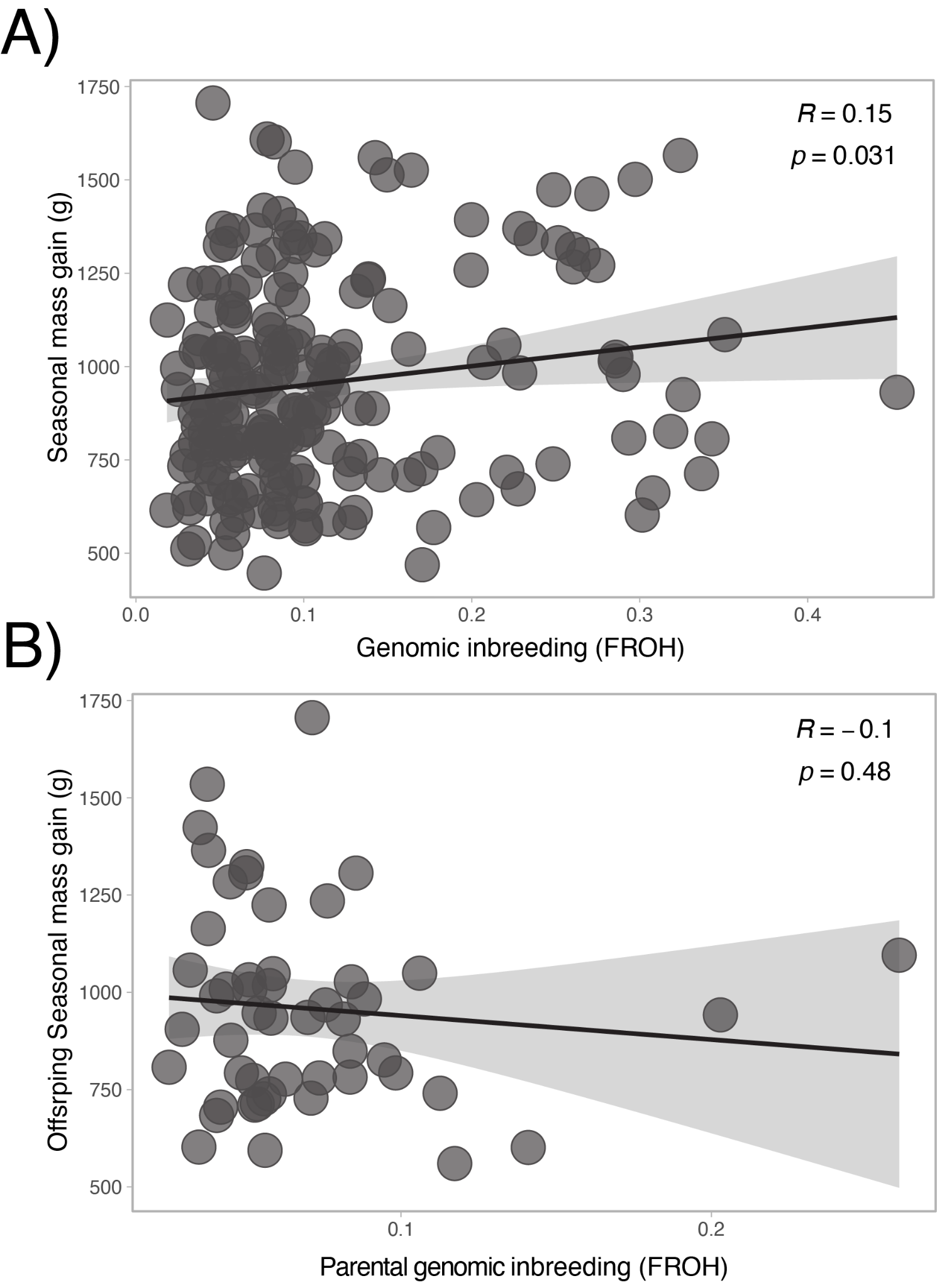
